# Genetic determinants that permit growth without the core septin Cdc12 in *Cryptococcus neoformans*

**DOI:** 10.64898/2025.12.17.694962

**Authors:** Piotr R Stempinski, Rooksana E. Noorai, Vijay Shankar, Lukasz Kozubowski

## Abstract

Septins are conserved cytoskeletal GTPases that regulate morphogenesis, cytokinesis, and cell-surface organization in fungi. In most model and pathogenic yeasts, including *Saccharomyces cerevisiae* and *Candida albicans*, core septin subunits such as Cdc3 and Cdc12 are essential and their loss results in lethal cytokinetic defects. In *Cryptococcus neoformans*, by contrast, deletion of *CDC12* yields viable cells that grow normally under permissive conditions, raising the question of how this pathogen maintains cell integrity without a functional septin complex. To address this, we compared the transcriptomes of wild-type cells and a *cdc12*Δ mutant. The *cdc12*Δ strain showed a focused set of transcriptional changes, with induction of genes involved in cell wall remodeling, stress-dependent morphogenetic programs, and pathways that promote fitness under compromised cytoskeletal conditions. Genetic analysis of the most highly induced genes uncovered factors that contribute to proliferation in the absence of Cdc12, including a previously uncharacterized aspartic-type endopeptidase that becomes essential when septin function is lost. These findings outline the compensatory circuitry that permits septin-independent growth in *C. neoformans* and help explain why this pathogen can tolerate septin loss. This intrinsic plasticity highlights *C. neoformans* as a powerful comparative model for studying the evolution and function of the septin complex.

## INTRODUCTION

*Cryptococcus neoformans* is an opportunistic yeast that causes life-threatening infections, particularly in individuals with weakened immune systems, such as patients with advanced HIV/AIDS, organ transplant recipients, and those receiving immunosuppressive therapies(Kwon-Chung et al., 2014; Lui et al., 2006; Perfect & Bicanic, 2015; Santos-Gandelman & Machado-Silva, 2019). Globally, *C. neoformans* is responsible for an estimated 150,000 cases of cryptococcal meningitis annually, leading to over 100,000 deaths, with the highest impact in sub-Saharan Africa(Kabir & Cunningham, 2022; Kwon-Chung et al., 2014; Lui et al., 2006). Beyond its medical relevance, *C. neoformans* has emerged as one of the most relevant model organisms for studying fungal pathogenesis, stress adaptation, and morphogenesis (Fernandes et al., 2022; Zaragoza, 2019). Its well-annotated genome, large number of genetic and molecular tools, and the availability of comprehensive mutant libraries have made it a key system for dissecting mechanisms of thermal tolerance, capsule biosynthesis, cellular division, and other virulence-related processes (Casadevall et al., 2000; Kong et al., 2017; Kronstad et al., 2012; L. Wang et al., 2012). *C. neoformans* is a basidiomycete fungus, evolutionarily distinct from the ascomycete model *Saccharomyces cerevisiae*, offering a valuable complementary perspective on fungal cell biology.

Septins are conserved cytoskeletal GTPases that play important roles in cell morphogenesis, cytokinesis, and cell surface organization (Douglas et al., 2005; Amy S Gladfelter et al., 2005). In yeast cells, septins assemble into hetero-oligomeric complexes that form filamentous structures at the mother–bud neck, serving as a scaffold for proteins involved in cell cycle regulation, membrane compartmentalization, and cell wall maintenance (Bridges & Gladfelter, 2015; Bridges et al., 2014; Douglas et al., 2005; Weirich et al., 2008). The septins Cdc3 and Cdc12 are essential components of the septin complex in *S. cerevisiae* and *C. albicans*, and their loss leads to lethal defects in cytokinesis (Flescher et al., 1993; McMurray et al., 2011; Warenda & Konopka, 2002). In contrast, cryptococcal cells lacking either Cdc3 or Cdc12 orthologs remain viable and proliferate relatively normally at optimal (non-stress) conditions (Kozubowski & Heitman, 2010). In our previous work, we reported that among approximately 80 temperature-sensitive single-gene deletion mutants, the strain lacking septin Cdc12 displayed the lowest maximal temperature (Tmax) at which cells remained viable and capable of proliferation (Stempinski et al., 2021). These findings suggest that the functional septin complex in *C. neoformans* is expendable for cellular maintenance and integrity under optimal environmental conditions but becomes essential for survival at elevated, mammalian host-relevant temperatures (Kozubowski & Heitman, 2010).

We hypothesized two non-exclusive scenarios that account for the viability of *C. neoformans* cells lacking Cdc12 under non-stress conditions. First, it is possible that septins are involved in stress response pathways. Second, it is plausible that the mutant that lacks Cdc12 has adapted to the absence of Cdc12 by transcriptional reprograming such that the selected cells (dominant marker selection) survive at non-stress conditions. To address the question of what mechanisms, account for the viability of *C. neoformans* cells lacking Cdc3 or Cdc12 at permissive temperatures, we performed RNA sequencing of WT and the *cdc12*Δ mutant. Our results reveal number of cell wall integrity and stress response related genes as upregulated in the absence of Cdc12 suggesting a mechanism responsible for septin-independent proliferation. Notably, this study has identified a novel gene, a putative aspartic-type endopeptidase whose function is essential for compensating for the loss of septin function in *C. neoformans* cells.

## MATERIALS AND METHODS

### Strains and Media

The wild-type H99 strain and the congenic *cdc12*Δ mutant strain of *C. neoformans* were used in the RNA sequencing experiment (Kozubowski & Heitman, 2010). Six deletion mutant strains were derived from the collection of single-gene deletion mutants of *C. neoformans* (Liu et al., 2008). Frozen cell stocks were kept at – 80°C and streaked onto YPD medium (1% yeast extract, 2% peptone, 2% agar, and 2% dextrose) and incubated at optimal room temperature (24°C).

#### Cryptococcus neoformans RNA extraction

Cell cultures were grown overnight (18 hours) at ∼24°C in 25 ml of YPD medium. 5 ml of overnight cultures were refreshed and incubated in 50 ml YPD for 3 to 4 hours to reach an OD600 of ∼1.0. The cultures were incubated in 25 ml YPD at room temperature (∼24°C). After final incubations, cells were harvested by centrifugation at 3000 x g for 5 min at 4°C. RNA extraction was performed with Bio-Rad Aurum Total RNA Mini Kit (Cat # 732-6820). Cells were mixed with 750 µl lysis buffer and approximately 50 µl glass beads (sigma G8772, 425-600µm) and mechanically homogenized at 4°C with Minibeadbeater (3450 RPM, Biospec Products Model 607) for eight 20-second beating cycles with two minutes rest in between. The supernatant with RNA was separated from the pellet by centrifugation (3000 x g for 3 min) and treated according to the manufacturer’s protocol. Each treatment and isolation was performed in three independent biological replicates.

### Transcriptomic analysis

Library preparation was completed using the Illumina TruSeq Stranded kit (Illumina, San Diego, CA, USA). Samples were sequenced to an average of 19.7 million reads per sample using an Illumina NextSeq 550 sequencing platform at 2×150 paired-end reads. Down-sampling was performed using seqtk v1.3-r106 (https://github.com/lh3/seqtk). Quality control was analyzed with FastQC v 0.11.6. Low-quality base sequences were trimmed using Trimmomatic v0.36. GSNAP v2018-07-04 software was used to count uniquely mapped reads that are aligned to reference genes. Statistical analysis for differential gene expression (DGE) was performed using edgeR v3.22.5. Analyzed genes were considered significantly differentially expressed if their false discovery rate (FDR) was greater than 0.05.

#### Cryptococcus neoformans mating

Single-gene deletion mutant strains (α), representing the group of genes with the highest upregulation at ∼24°C in *C. neoformans* lacking Cdc12, were mated with a strain (a) carrying a *CDC12* deletion (Kozubowski & Heitman, 2010). To induce mating, cultures of two different mating types (MATα and MATa) were grown overnight in YPD liquid at ∼24°C, washed with PBS, mixed in a 1:1 ratio, and spotted onto MS mating media. Colonies were incubated in the dark at ∼24°C for 10-12 days (Lin et al., 2008; Sun et al., 2019; Xue et al., 2007). For each strain, 32 spores, originating from at least four different basidia, were transferred onto the YPD semisolid media with SporePlay Tetrad Dissection Microscope (Singer Instruments). To confirm the results of the mating and identify strains with double gene deletion, colonies were tested for growth on YPD media supplemented with a selection drug (NAT and/or G418).

### Serial spot dilution assay

Overnight cryptococcal cell cultures were washed twice with DPBS, adjusted to a final concentration of 3.3× 104 cells/μL, and used for a 10-fold serial dilution. 3 μL of cell cultures were spotted onto YPD media or YPD media supplemented with SDS (0.01%), Fluconazole (8 µg/ml), or Congo Red (5 mg/ml). Plates were imaged after the cells were incubated at ∼24°C for three days.

### Gene identification and GO analysis

Genes of *C. neoformans* identified in this study were analyzed using the FungiDB database (https://fungidb.org/). Functional analysis of the selected groups of genes was performed on the differentially expressed genes to identify overrepresented Gene Ontology (GO) molecular functions and biological processes.

### Protein sequence analysis

Amino acid similarity analysis was performed using the protein-protein BLAST (NIH). Analysis of the secondary and tertiary structure of the protein encoded by the CNAG_05305 gene was performed using the AlphaFold 3 tool (Abramson et al., 2024). Computational protein localization prediction was analyzed using PSORT: Protein Subcellular Localization Prediction Tool (https://www.genscript.com/psort.html)

## RESULTS

### Loss of septin Cdc12 induces cell wall remodeling and stress response pathways in *C. neoformans*

In *Saccharomyces cerevisiae*, deletion of the septin Cdc12 is lethal due to its essential role in septin complex assembly and cytokinesis (Bridges et al., 2014; Amy S Gladfelter et al., 2002; A S Gladfelter et al., 2001; McMurray et al., 2011). In contrast, while loss of Cdc12 in *Cryptococcus neoformans* disrupts septin ring formation at the mother–bud neck, *cdc12*Δ cells remain viable at temperatures below ∼32°C (Kozubowski & Heitman, 2010; Stempinski et al., 2021). We hypothesized that *C. neoformans* adapts to the absence of Cdc12 through a supporting gene expression pattern that allows for growth under optimal conditions. To test this hypothesis, we analyzed and compared transcriptomic profiles of the *cdc12*Δ mutant and wild-type (H99) strains grown in rich medium (YPD) at ∼24°C. RNA sequencing revealed extensive transcriptional changes in the absence of Cdc12. A total of 134 genes were differentially expressed between the *cdc12*Δ mutant and wild type (FDR < 0.05). Strikingly, 131 genes were upregulated, while only three genes encoding proteins with unknown function were significantly downregulated (CNAG_06347, CNAG_04756, CNAG_00399), (Supplementary Table 1). All genes with logFC> 2 were listed in Table 1.

**Table 1.** Table with the list of 23 genes with the highest level of upregulation (>2 logFC) in the *cdc12*Δ mutant in comparison to wt strain of *C. neoformans*.

| Gene ID | logFC | Gene Name | Product Description | Protein size |
| --- | --- | --- | --- | --- |
| CNAG_05305 | 5.4 |  | hypothetical protein | 608 aa |
| CNAG_04891 | 4.2 | Ril1 | Ricin B lectin domain-containing protein | 204 aa |
| CNAG_03833 | 3.6 |  | hypothetical protein | 418 aa |
| CNAG_00587 | 3.3 | Ril3 | Ricin B lectin domain-containing protein | 185 aa |
| CNAG_07399 | 3.3 |  | hypothetical protein | 122 aa |
| CNAG_02339 | 3.1 |  | DUF1275 domain protein | 311 aa |
| CNAG_02381 | 3 |  | DH domain-containing protein | 479 aa |
| CNAG_05075 | 2.9 |  | solute carrier family 20 | 591 aa |
| CNAG_03650 | 2.8 |  | hypothetical protein | 67 aa |
| CNAG_02544 | 2.6 | SWI5/SAE3 | DNA repair protein | 102 aa |
| CNAG_05258 | 2.5 | SMG1 | glucose-methanol-choline oxidoreductase | 588 aa |
| CNAG_03630 | 2.5 |  | hypothetical protein | 246 aa |
| CNAG_01653 | 2.5 | CIG1 | cytokine inducing-glycoprotein | 281 aa |
| CNAG_07969 | 2.4 |  | hypothetical protein | 309 aa |
| CNAG_00323 | 2.3 |  | Kre9/Knh1-like domain containing protein | 167 aa |
| CNAG_00177 | 2.3 |  | hypothetical protein | 530 aa |
| CNAG_00795 | 2.3 | CFL1 | Putative flocculin | 309 aa |
| CNAG_04556 | 2.2 | SEC62 | translocation protein | 312 aa |
| CNAG_05458 | 2.2 |  | endo-1,3(4)-beta-glucanase | 368 aa |
| CNAG_07877 | 2.2 |  | hypothetical protein | 175 aa |
| CNAG_06715 | 2.1 |  | hypothetical protein | 198 aa |
| CNAG_01994 | 2.1 |  | SET domain-containing protein | 1006 aa |
| CNAG_06027 | 2.1 |  | aryl-alcohol dehydrogenase | 400 aa |

Among the upregulated genes, 55 (42%) were either previously characterized or could be inferred by sequence homology. Notably, several genes associated with cell wall biosynthesis and remodeling were strongly induced in the *cdc12*Δ mutant, including *CHS5* and *CHS7*, encoding chitin synthases (CNAG_05818 and CNAG_02217), *AGN1* (CNAG_07736), which encodes a glucan endo-1,3-alpha-glucosidase, and three glucanases (*CNAG_05458*, *CNAG_06336*, *EXG104* / *CNAG_02225*), as well as a glycosyl hydrolase (*CEL1*, CNAG_00601)(Banks et al., 2005; Donlin et al., 2014; Mukaremera, 2023; Probst et al., 2023; Reese et al., 2007; Rodrigues et al., 2018). The upregulation of these genes suggests that reinforcement or remodeling of the cell wall may compensate for the loss of structural integrity caused by the absence of a functional septin scaffold (Figure 1). Interestingly, two of the most highly induced genes in the *cdc12*Δ mutant encode proteins containing a Ricin B lectin domain (*CNAG_04891* and *CNAG_00587*). Knowing that Ricin B lectin domains are carbohydrate-binding modules that interact with polysaccharides, their upregulation may indicate a supporting mechanism contributing to capsule organization or cell wall maintenance in septin-deficient mutant cells (De Coninck et al., 2024; Xu et al., 2022).

**Figure 1.**
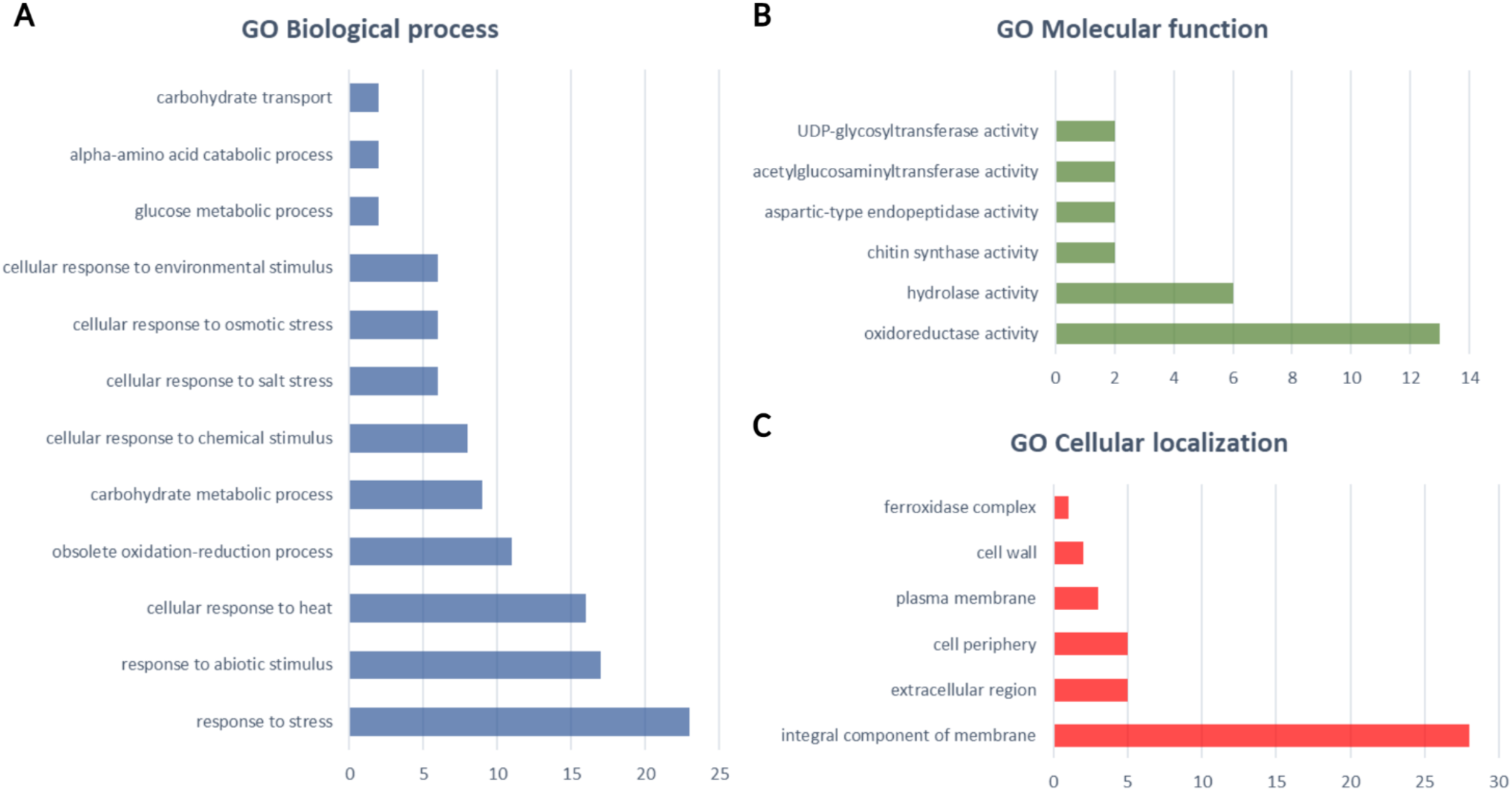
Bar graphs representing GO terms analysis for (A) biological processes, (B) Molecular function and (C) Cellular localization of *C. neoformans* genes upregulated in the *cdc12*Δ mutant. The X-axis represents the number of genes in each GO term.

In addition to genes associated with cell wall and surface architecture, we observed upregulation of two components of the iron acquisition system essential for cryptococcal virulence: the cytokine-inducing glycoprotein *CIG1* (CNAG_01653) and the ferric transporter *CFT1* (CNAG_06242)(Cadieux et al., 2013; Garcia-Santamarina et al., 2017; Han et al., 2012; Jung et al., 2008). Furthermore, two genes encoding a glutathione S-transferase (*CNAG_01874* and *CNAG_04110*) involved in glutathione metabolism and reactive oxygen species homeostasis were also significantly upregulated, suggesting activation of oxidative stress response pathways (Missall et al., 2005; Wangsanut & Pongpom, 2022). Finally, *CFL1*, encoding a putative flocculin, showed one of the highest levels of induced expression (L. Wang et al., 2012, 2013). *CFL1* is known to be a hypha-specific adhesin and among the most strongly expressed genes during sexual development, suggesting activation of another mechanism supporting cellular morphology and cellular growth in the absence of the septin complex. (L. Wang et al., 2012, 2013).

We next performed Gene Ontology (GO) term enrichment analysis to characterize the biological processes, molecular functions, and subcellular localization of the proteins encoded by genes differentially expressed in the *cdc12Δ* mutant strain. The analysis of biological processes showed strong enrichment for genes involved in responses to stress, abiotic stimuli, and heat stress (Figure 1A). Additional categories included genes involved in the oxidation-reduction process and carbohydrate metabolism. GO analysis of molecular function further highlighted that the dominant protein groups are engaged in oxidoreductase and hydrolase activities (Figure 1B). Finally, GO analysis of cellular localization indicated that most proteins are integral membrane proteins and are localized to peripheral and extracellular regions of the cryptococcal cell (Figure 1C).

Collectively, these results demonstrate that the absence of Cdc12 septin in *C. neoformans* elicits significant alterations in gene expression. Transcriptional adaptation is centered on the expression of genes involved in cell wall synthesis, oxidative stress response, and nutrient acquisition. This adaptation likely enables *C. neoformans* cells to maintain cellular integrity and viability in optimal environmental conditions, despite the loss of a key septin component that is essential in most other fungal species.

### Phenotypic analysis of strains lacking genes that were highly upregulated in the absence of septin Cdc12

To explore the nature of the putative suppression of a potentially deleterious phenotype in the absence of the septin complex, we selected six mutant strains carrying deletions of the most upregulated genes in the *cdc12Δ* mutant strain. The protein sequences of all six highly upregulated candidates were analyzed with BLASTP (https://blast.ncbi.nlm.nih.gov) and FungiDB (https://fungidb.org/) for potential homologous proteins in other fungal species. Interestingly, homologs of these genes were found only in the order Tremellales, which includes the genera *Cryptococcus* and *Tremella*. Serial dilutions of WT, *cdc12*Δ, and those six selected mutant strains were spotted onto YPD media supplemented with the cell wall-disrupting agent congo red, or a subinhibitory concentration of the plasma membrane-disrupting antifungal drug fluconazole and incubated at 30°C or 37°C (Figure 2). Consistent with previous observations, incubation on solid YPD at 37°C confirmed that *cdc12Δ* was extremely sensitive to elevated temperature, whereas all other deletion mutants and the WT strain grew well. Interestingly, deletion of gene CNAG_05305 resulted in a drastic growth defect on YPD supplemented with 8 µg/mL fluconazole, exceeding previously reported sensitivity of *cdc12Δ* mutant cells (Kozubowski & Heitman, 2010; Stempinski et al., 2021). This finding suggests that CNAG_05305 plays a role in plasma membrane homeostasis and/or stress response pathways that are critical for cell vitality in the absence of a functional septin complex. In contrast, supplementation with 8 µg/mL congo red produced mild growth inhibition of the *cdc12Δ* strain and had no visible impact on the other mutants. These results suggest that neither of the six analyzed genes is essential under cell wall stress conditions. Furthermore, among the six genes, only the CNAG_05305 was essential under conditions that compromised plasma membrane.

**Figure 2.**
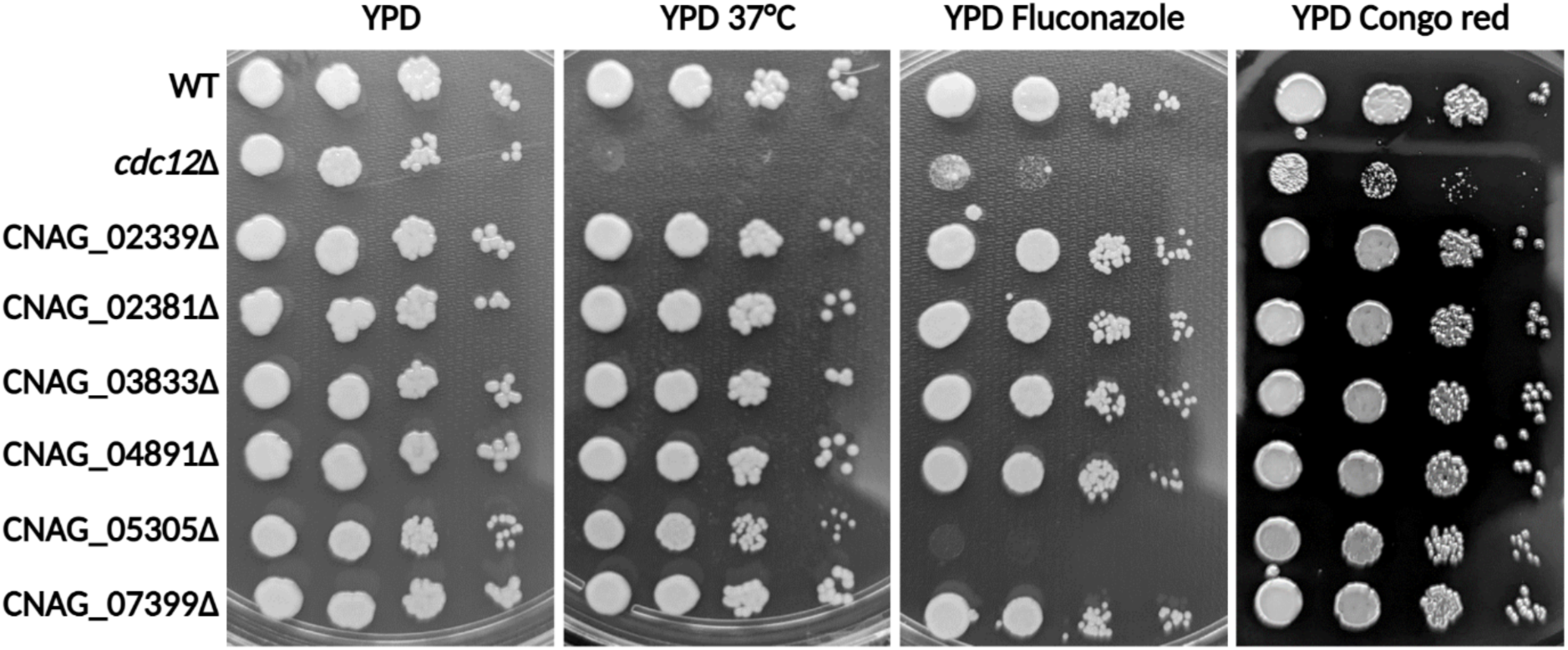
Spot assay of WT, *cdc12*Δ, and selected mutant strains. Threefold serial dilutions of each cryptococcal strain were spotted onto YPD media, YPD media with 8 µg/ml fluconazole, and YPD media with 5 µg/ml congo red. Plates were incubated at 30°C or at 37°C as indicated.

### Genetic interaction of *CDC12* and putative Aspartic peptidase highlights compensatory mechanisms important for morphogenesis

The septin complex plays a vital role in regulating the cell cycle and cell wall synthesis in *S. cerevisiae* (Bertin et al., 2008; Douglas et al., 2005; Amy S Gladfelter et al., 2002; A S Gladfelter et al., 2001; Glomb & Gronemeyer, 2016; McMurray et al., 2011; Momany & Talbot, 2017). *C. neoformans* cells lacking a functional septin complex display aberrant morphology of the hyphae, increased sensitivity to cell wall-damaging agents, and significantly decreased virulence (Kozubowski & Heitman, 2010). Loss of septin function is predicted to produce pleiotropic morphological defects, including cytokinesis abnormalities. Interestingly, at temperatures below ∼32°C, the *cdc12Δ* mutant maintains largely normal morphology, likely due to a supportive gene expression profile.

As high expression of a gene in the absence of Cdc12 suggests this gene is essential in the absence of this septin, we hypothesized that eliminating both *CDC12* and such a gene would lead to a synthetic lethal or synthetic sick phenotype in the double mutant. To confirm the supporting role of selected genes in the septin-deficient strain of *C. neoformans*, we performed mating between the *cdc12*Δ mutant and deletion mutants corresponding to the group of most strongly upregulated genes in the *cdc12*Δ background. Among all tested progeny, only the double deletion of *CDC12* and CNAG_05305 led to progeny with an apparent growth defect (Figure 3).

**Figure 3.**
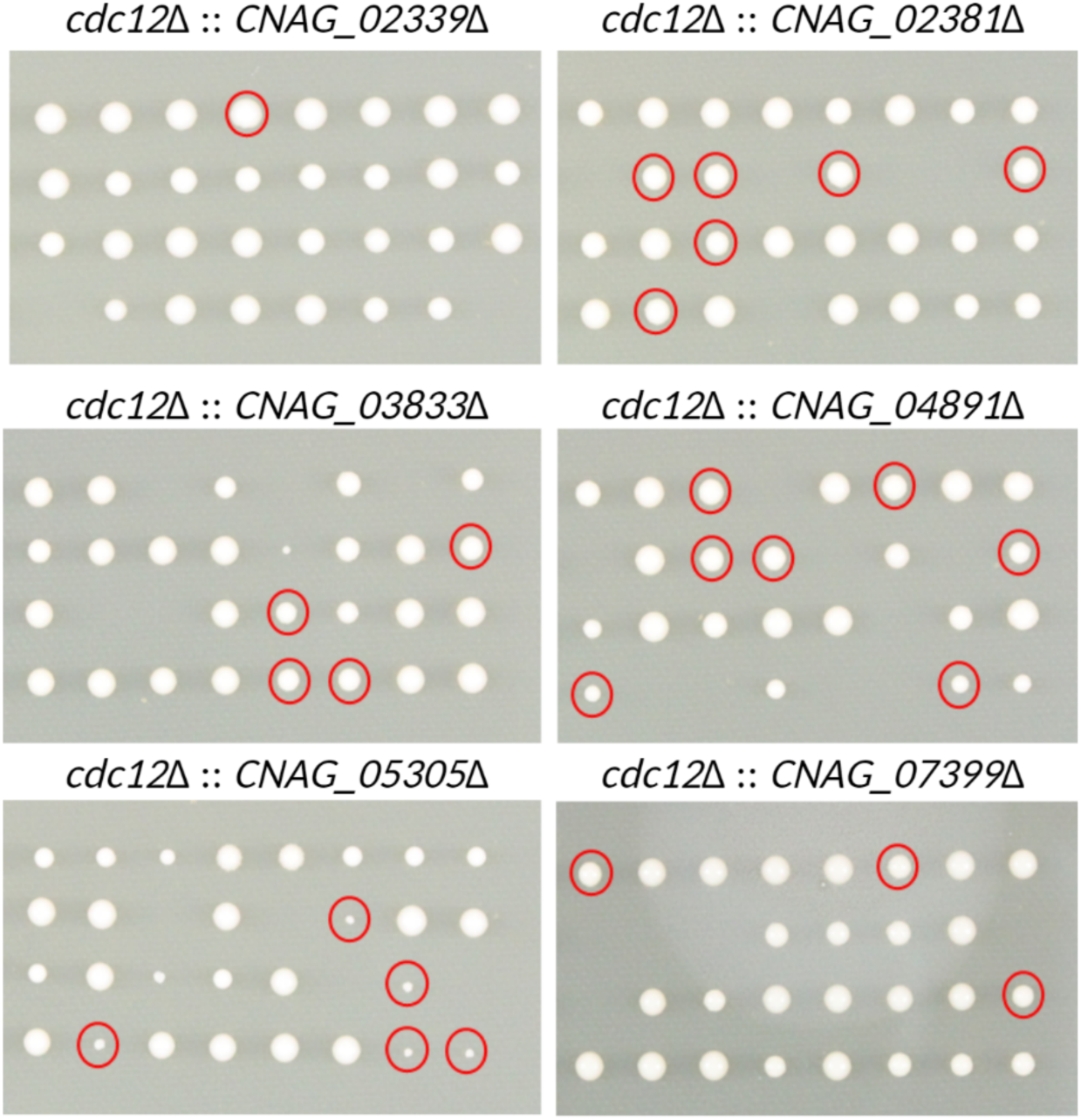
Qualitative growth analysis of single spore colonies. Spores obtained by mating of the *cdc12*Δ mutant with deletion mutants of the genes with the highest upregulation in the septin *cdc12*Δ background. Red circles indicate confirmed colonies of double-deletion mutant strains.

### Structural prediction and functional analysis of CNAG_05305, a putative Aspartic peptidase with a CCHC-type zinc finger motif sequence

Initial *in silico* analysis of the cryptococcal protein encoded by CNAG_05305 using the AlphaFold 3 Protein Structure Database identified it as a protein of unknown function with a predicted molecular mass of approximately 68.1 kDa and a tertiary structure comprising three distinct protein domains (Figure 4A and 4B). The amino acid sequence analysis using InterProScan indicated the presence of an aspartic-type endopeptidase domain and a CCHC-type zinc finger motif domain, which is typically associated with nucleic acid binding (Figure 4C)(Benhalevy et al., 2017; Mackeh et al., 2018; Y. Wang et al., 2021). Detailed analysis of the indicated CCHC-type zinc finger-like motif revealed the existence of a 13-residue domain with the CX2CX3HX4C consensus sequence (<u>C</u>FT<u>C</u>GAY<u>H</u>WSDQ<u>C</u>). Further evaluation of the zinc finger-like domain in the protein sequence of homologous proteins in other, closely related fungal species confirmed the highly conserved character of this domain (Figure 4D and Supplementary Figure 1). Subcellular localization analysis using the WoLF PSORT tool predicted that the protein encoded by CNAG_05305 is most likely localized to the nucleus or the cytoplasm, further supporting a regulatory or processing role for this protein.

**Figure 4.**
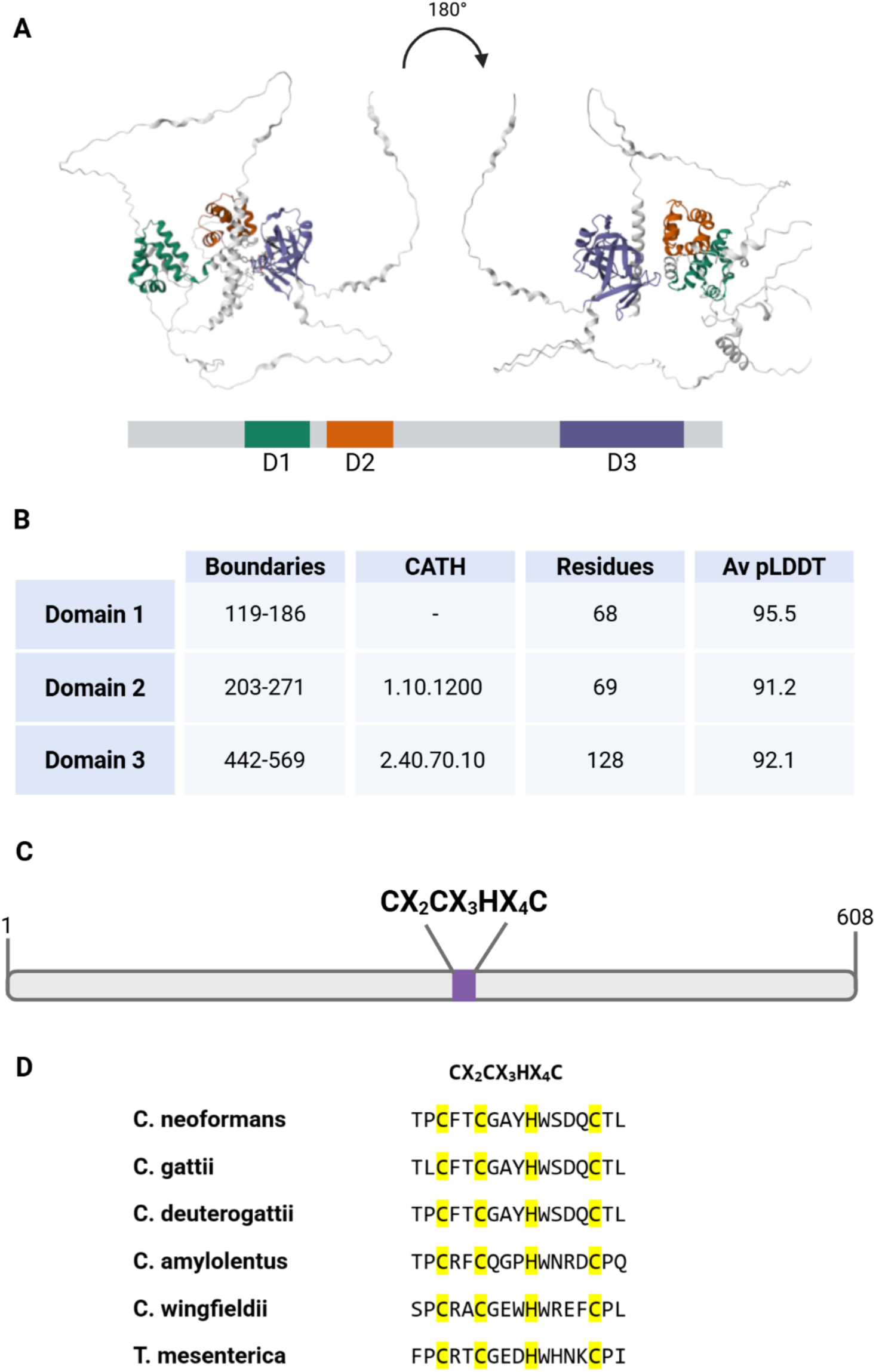
Predicted structure and domain organization of the CNAG_05305 protein product, a putative aspartic peptidase. (A) Predicted three-dimensional structure of the CNAG_05305-encoded protein generated using AlphaFold3. Two orientations of the model, rotated 180° along the x-axis, are shown to illustrate the overall protein architecture. The three predicted structural domains identified by AlphaFold are highlighted in distinct colors and labeled as domains. (B) Summary table of the structural features of the CNAG_05305-encoded protein, indicating the boundaries of each domain, corresponding CATH classification, number of residues, and average predicted local distance difference test (pLDDT) score. (C) Visualization of the putative zinc finger motif, represented by a 13-residue region containing the conserved CX₂CX₃HX₄C sequence within the protein. (D) Multiple sequence alignment of the CX₂CX₃HX₄C zinc finger region from several *Cryptococcus* and *Tremella* species. Conserved cysteine and histidine residues are highlighted in yellow.

Further microscopical analysis confirmed previously published observations that most *cdc12*Δ cells grown in YPD at ∼24°C resembled wild-type cryptococcal cells, with only a minor fraction of the population presenting elongated or misshapen forms. Similarly, the single deletion of CNAG_05305 resulted in predominantly normal cellular morphology, with a very minor number of elongated cells. In contrast, the double-deletion mutant strain showed severe morphological defects across the entire cell population, highlighting an important role for CNAG_05305 in maintaining proper cell shape and integrity when the septin complex is compromised (Figure 5).

**Figure 5.**
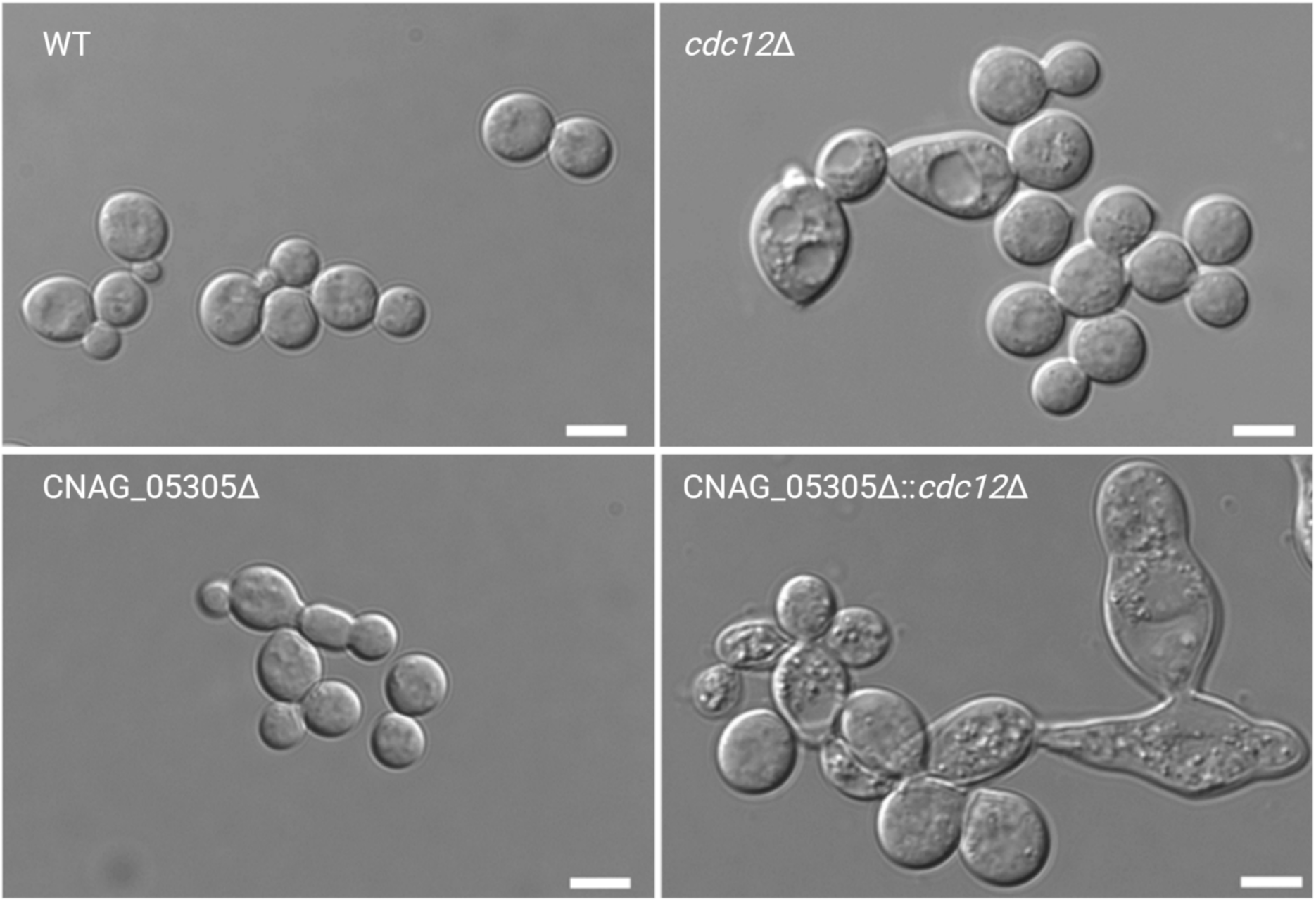
Cell morphology of the WT and deletion mutant strains. The *cdc12*Δ mutant and the *cdc12*Δ, CNAG_05305 double deletion mutant presents abnormal cell morphology. Strains were grown for 24 h in liquid YPD at 24°C and cellular morphology was examined. Scale bars represent 5 µm.

## DISCUSSION

The main goal of this study was to investigate the role of the septin complex, specifically septin Cdc12, in the cellular physiology of *Cryptococcus neoformans* and the putative supporting molecular mechanisms that allow the fungus to compensate for the lack of Cdc12 under “optimal” growth conditions. Transcriptomic analysis revealed that the *cdc12*Δ mutant requires a major transcriptional adaptation, characterized by the increased expression of multiple genes involved in cell wall biosynthesis and integrity, oxidative stress response, and nutrient acquisition. Identification of genes encoding membrane-associated proteins and extracellular components suggests that septin deficiency impairs cell surface integrity. Functional validation of the protein encoded by the gene presenting the highest level of upregulation, CNAG_05305, a previously uncharacterized Asp protease-containing protein, confirmed it as a critical factor for survival under fluconazole-induced stress. Double deletion of *CDC12* and the CNAG_05305 resulted in synthetic growth defects and severe morphological abnormalities, providing direct evidence of genetic interaction and a compensatory role for the CNAG_05305-encoded putative Aspartic peptidase in maintaining cellular morphogenesis in the absence of Cdc12.

Comparison of the septin complex biology in *S. cerevisiae* and other fungi emphasizes its conserved and versatile functions. In *S. cerevisiae*, Cdc12 is essential for septin complex assembly, cytokinesis, and cell survival (Bridges & Gladfelter, 2015; Bridges et al., 2014; Amy S Gladfelter et al., 2005). In contrast, *C. neoformans* cells can tolerate the absence of the septin complex under optimal growth conditions, reflecting species-specific differences in septin network robustness and adaptive capacity. Similar temperature-sensitive phenotypes have been reported in other pathogenic fungi, suggesting that septin-dependent processes are particularly critical under host-relevant stress conditions.

CNAG_05305 may support *C. neoformans* cells lacking septin complex via several potential mechanisms. The analysis of protein sequence revealed that CNAG_05305 encodes a putative aspartic-type endopeptidase with a CCHC-type zinc finger motif, features often associated with RNA metabolism and general transcriptional regulation (Benhalevy et al., 2017; Mackeh et al., 2018; Y. Wang et al., 2021). The cytoplasmic or nuclear localization of this protein suggests that its function may involve regulating stress-related transcription or mRNA dynamics important for indirectly reinforcing the plasma membrane or cell wall. The abnormal phenotype observed in the double-deletion mutant of *CDC12* and CNAG_05305 supports the idea that these proteins operate in parallel, partially overlapping pathways that together maintain cellular organization and morphogenesis. This observation is further supported by the fact that in the *cdc12*Δ strain, CNAG_05305, present only in closely related fungal species, showed the highest level of upregulation among all detected genes (>5 logFC).

Together, these findings reveal that *C. neoformans* adapts to the absence of a key septin by unique transcriptional reprogramming and activation of genes, such as CNAG_05305, which are uniquely present in the genomes of fungi that belong to the order Tremellales. This increased adaptive capacity, may explain why elimination of the septin complex in *C. neoformans* is not lethal in contrast to *S. cerevisiae* or *C. albicans* (Flescher et al., 1993; Frazier et al., 1998; Warenda & Konopka, 2002). Future studies will shed more light on the specific role of the CNAG_05305 in *C. neoformans* morphogenesis and the mechanism of its contribution to stress response and antifungal resistance.

