## Supplemental Figure 1 for "Genetic determinants that permit growth without the core septin Cdc12 in *Cryptococcus neoformans*"

***Cryptococcus neoformans* H99**

>XP_012048840.1 hypothetical protein CNAG_05305 [Cryptococcus neoformans H99]

MNNRLATHLQHVDTIYQQRFNELHDLFQATHGATLRHSTPLHQADPSLDTSSSTSPTPPLRPTSPPTCPP

TQPPTIPLDPPSSTSVIGKNDSVLSTSQPTRLKASDLPKFYGRDDDDIVYWVQKISSIKRGSQATDSDIL

RLLPSLLRDNAFKYYARLPEKEHTSLTTWAAWSKELQDRFLPPNSLDDLKEKCIRRYLGPNETFANYYED

KVYLQQFLFPANTEDSLLVRDILGGIPATLRAQLQTGVTPTMSLREFRLHVLQIEPRFHPQLFPTSHRHP

ADRTRTQLNNHSNTSFANSSIARSSRFLPGNRPEKPSTPCFTCGAYHWSDQCTLKKRQQQQYNNSRPSFT

TNNPATSTSSDGRSSFPSWARPPSNTNSPNYKPPANSVNSIANKDGYRFDSPKPSIHTLCHSTFPRPSAH

LPTHALPQLQLHRTHLNKTPAYATVKFNSNNPSATTHRAVIDTGASLSAISQEYANKHLSHLPRHPFTNF

QLNGIGKSSSAGYHLVDIHFPTKEGYHIVIPTALFVIDNISVDLILGLDFLLPHSAKIDLRDGILSFPRH

SGTILLSCLSSALPFPPIPLPRPPRRFIAKSSRVQTMLFPDPVAERTI

***Cryptococcus gattii* WM276**

>XP_003192864.1 Hypothetical Protein CGB_C5480W [Cryptococcus gattii WM276]

MSTSPSTRPRALSVGADARSQDSIFPPPSIVSPRRMSITPQPKPLAHEIAHHYEVEIENRVTTCLKRIEE

VHQQRLNELNTLMQSMSDMMPTVSSHSIPLDHIVSSPNTSLPSLPIPPVPLRHTLPSQPTVTSTPTQTSA

QPTRLKASDLPKFYGRDDDDIVEWVQKVGSLKRVFEATDRDILRLLPSLLHELQDRFLPPNRLDDLIYRC

IRRYLGQNEKFVTYYEDKVYLQQFLFPANTEDFLLIRDILSGILSTLRAQLQTAVTPNMSLCDFRCHVLQ

IEPGFRPHLFPSSHRHPADRARHQRNNHPQRSSDTSTSARPFRHTPGTRPDKPCTLCFTCGAYHWSDQCT

LKRQQHQQYGARPPPATNNPGSSTSSTSRRFFLSWARPASNTTSHSYQPLSNSVNSIQPKDNNRFNSPKP

SINTLLHSHSPAPKPLHALPPLQLQQKHLNMTPAFANIKFNSNHPSALTHRAVIDTAPGHWQILLSRISP

G

***Cryptococcus deuterogattii* CA1014**

>KIR75865.1 hypothetical protein I310_00563 [Cryptococcus deuterogattii CA1014]

MYITMRHPVPTPCFTCGAYHWSDQCTLKQQQQQQHNATWPSSNANNPATSTSLNSCSSFPSWACPSPNT
N

SSNYKPPANLVNSIANKDGYCFNSPKPSVHTLLHSTPTCSSTHSPVHAIPQLQLQQTHLNKTPAFATVKF

NSNHPFASTHHAIINTGASLSVIRQEYAQKHLSHLLCHPFKNFQLNGIGQSSSTGYHLIDIHFLTKEGYN

TIILTALFIINNISVNLILGLDFLLPHHAKIDLQDGILSFPCHPGTILLSCQSGPLIPPHLPPLPSPIPC

PFVPSIHVKEAFTILPVHQAHIETTIDVLPHTPQYLVSPTSTGKYLHVACCVGSSTASKHFVLVLNTGDR

PIPVSGGHLLGYPLPLHTACPTPAVHTINTVAIPETDSDAEPFPITDLKINPDLTPEQSEALHQVLMEQC

HIFGFGSQQLGSTNLATMNVDTGNHPPISTPPYRISPQGHATIDEKLTELLNQGIIEEADSPWASPAILV

***Cryptococcus amylolentus* CBS 6039**

>XP_018994010.1 hypothetical protein L202_04479 [Cryptococcus amylolentus CBS 6039]

MAQPATQPSRRNLRSQGDALLDGPADVVTNQPQPTPTLPSGPSLQSEVDIRPSNFQPPPSIDDTPTSEHH

ITTHQDLVHGPAPTVGYLMHQNQSLLSSLAEMRDEMRSLRLSQSRDTTPVSPPSRPHAPKLKHEMFPKFR

GQDNEDVDTWVTSVTAIFEFSGAADTDLLLLLPALLQGSAQRWFTNLEPERRRSLCTWSQWSEAIRSVYL

GANHASRLRNKCIFRRLKTTDGLSKDIYPAGTPDVDRIDDIINGLPEHMRKLARIDAQLHPNLDDFRRAL

IDGESSMRPELKQNRRRQDKAPPTSQNKHQQRDDKNSSSTLSSPRTPCRFCQGPHWNRDCPQQATAPPRP

TQTMAQPNPSPSTAPTYGNFRLASRPDSDRPLGFRPAPASANGTPLGSRSSSVETTSGPIKSITVSGAVS

LGSTVAELGTETVGGVDTWNARGARERGGDSRNVGRGGSLSLGIRDSRSLVGGGSLSLGIGVSRSLSRGR

RSGPYEHGVSTIWLYTCHCTANSSGKKVESPRKPPQTHEQQTTLSCFYGNESW

***Cryptococcus wingfieldii* CBS 7118**

>XP_019029938.1 hypothetical protein L198_06153 [Cryptococcus wingfieldii CBS 7118]

MPPATRSGPSSAPVPSDADTPTSSLTSLADPVETGGPSQAQSQADPSSEVSNLGDAVVPQTLPAEDGTAT

DQSQVASGSRATDHPTDLPARGTSLAPSETPSSWHSDLQVLSQNQALLTRLLHTMHEQNILRAQSLPTSQ

TPPPPDNPSSQPSFERRRRLRPADLTKFGGSDTEDVDMWLEKLTAALEHADYPESELLSNLPFLLEGKAL

DWFTDLGPVRRDYQTWDEWRVVFKNAFRIPDFEGVMRRKCIARRLQPFESFADYFDDKRRLQRWVYPVGT

SSKDLITDIVEGIPLVMRALIKASTPPGASLDDFRRIMLDLQPSLRSQFPTPNDKPPRVHNDDQQDRSRT

APSRQSPQTRAAAPPSPCRACGEWHWREFCPLNSPRPFNTPPSDSYGRSQNGFDDHQRHDSQSRTFAGSG

SNGISRPSDNGYQDQQDTRPFQSHSPGNV

***Tremella mesenterica***

>RXK37002.1 hypothetical protein M231_05709 [Tremella mesenterica]

MSTPNIPSTDQGLANPGTNTHTQVTDDHIRTLCSQLLNHHNTNIDNQIVQLQNAIDHLESTHAQEIDDLL

SQLNDLQEKCDNLQHQLDTTTNISVPTTNHSTVPLVNSTSHIKLKVTELPKFTANSTSESVDDWVNKITA

ILQQSGAPDTEITAKIPLLLQFAALDWFNTLSNIDRAKCKTWADWSTLFKSTFRAANYSAKLLALARDRH

LLPEESVTTYFWQKVKLLRDVHGSLITNQVLVNEILAGLPTSMSTFLNVNPNIELTEFHRLLVDKEDGLR

TLGFTPNFSTNLDQDQDTTNIFDNNYDSEEDVLDNYANNTNTIPTAPPFPCRTCGEDHWHNKCPIRNGVY

HDNNTIDHSNHDNADTYSHHDPAAPDNNSHNWGHHQNTCTSNKTIYKDNPNLQPITKPRQWVPLNILFCL

ETSSDTQLSGSQTPQAHQQAIGLVRKMGSFFSHRRASSFPLPWSSSQDESIFSTSTYEPPSYQNDNGSFN

IIQSKLFSRTRDHGTGESTMWGKIKSWFRLNSSVSYSNPTIPPELYVTLAESLTRLQLHKTLLNFSLSSK

SHYRLISPVLYREAILPLNDNVVDKLGDFYLHYKIFPQSRDSLIDPDFSFLTWDPNYLASSTSARLAWRC

QWARTVRIRTSPQTLQSLSMTSIVQTLSKIELCLFPRLTKLSLRRSASRKSATLLRLLALKSEIDHLLLF

LGSSHLDIHFPWFRDYLTLHDPVHIDRMVVRVDIKKNKWYFYPVKLSPTTGDDRNVSKDDFERQFDAKKS

DRDTLPRFHILSSPPKDRRRSTVASLNHYLKENTNEAGNFAKILTLGSVDLDPCIVFDPDKQKAIICRKI

RIVKKEVGQ

| | **$CX_2CX_3HX_4C$** |
|---|---|
| *C. neoformans* | TPCFTCGAYHWSDQCTL |
| *C. gattii* | TLCFTCGAYHWSDQCTL |
| *C. deuterogattii* | TPCFTCGAYHWSDQCTL |
| *C. amylolentus* | TPCRFCQGPHWNRDCPQ |
| *C. wingfieldii* | SPCRACGEWHWREFCPL |
| *T. mesenterica* | FPCRTCGEDHWHNKCPI |
